# How Public Transport affected the Propagation of Zika and Microcephaly within Rio de Janeiro early in 2015

**DOI:** 10.1101/345454

**Authors:** M. Armstrong, F.C. Coelho, L. Bastos, V. Saraceni, C. Lemos, M. Silva, L. Santana, J Crespo, M. Watanabe

## Abstract

From mid-2015 to the end of January 2016, 47 cases of microcephaly were observed in the city of Rio de Janeiro, that were not due to other viral infections (syphilis, toxoplasmosis, herpes & cytomegalovirus). These children were conceived from Dec 2014 to April 2015, far too early to be explained by the officially recorded cases from October 2015 onward. Zika must have been rampant in the city from late 2014 onward. In the first half of the paper we study how the geographic spread of microcephaly cases evolved from mid-2015 to January 2016 (and hence Zika 6-9 months earlier). Cases were not evenly spread in proportion to the number of births; they were preferentially located in the northern suburbs apparently following the public transport routes, with virtually no cases in favelas and none in the southern suburbs (Zona Sul). One key difference between the transport systems in the northern and southern suburbs is that the metro & rail system in the north is above ground in the north whereas in the southern part the metro is underground with air-conditioning in carriages and forced ventilation on the platforms. The train system does not extend to Zona Sul.

In the second half of the paper we postulate that the air-conditioning and ventilation prevent mosquitos from biting people who are waiting on platforms in Zona Sul. Agent-based simulations are used to test this hypothesis. After confirming this, we postulate that providing air-conditioning and/or forced ventilation on the rail-metro transport hub in the city center (Centro) would significantly delay the propagation of arboviruses in the city, possibly preventing epidemics. One advantage of this proposal is that it does not require the use of insecticides.

## Introduction

In October 2015, the Brazilian Ministry of Health (MoH) notified the World Health Organization [^1^ http://www.who.int/csr/don/20-november-2015-microcephaly/en/] of an unusual increase in the number of microcephaly cases among newborns in the state of Pernambuco, northeastern Brazil. On 11 November, the MoH declared a national public health emergency to give greater flexibility to the investigations and ten days later, the WHO released a Disease Outbreak News on this (Heymann et al, 2016). On 28 November the MoH [^2^ http://portalsaude.saude.gov.br/index.php/cidadao/principal/agencia-saude/21014-ministerio-da-saudeconfirma-relacao-entre-virus-zika-e-microcefalia] officially announced that there was a link between microcephaly and the Zika virus. It took several more months before the link between the Zika virus and microcephaly was recognized in other countries. In the USA, the CDC only released a statement to this effect on 13 April 2016 [^3^ http://www.cdc.gov/media/releases/2016/s0413-zika-microcephaly.html].

Rio de Janeiro also experienced an increase in the number of cases of microcephaly and of Zika. From 25 October 2015 onward it was compulsory to notify the Health Authorities of any cases of Zika but before that, the virus was probably present in the city but unnoticed. This paper focuses on the early stages of the epidemic. The questions that we address are: were the microcephaly cases evenly spread throughout city, and how was the Zika virus propagated within the city? The microcephaly cases turned out to have been concentrated in certain parts of the city but absent from others. Based on the spatial distribution of these cases, we postulate that the transport system played an important role in this transmission: the above ground rail and metro lines contributed to the propagation but that the underground metro lines slowed it down because of the forced ventilation on platforms and the airconditioning in carriages. We argue that increasing the ventilation on train platforms and in waiting areas at transport hubs would have significantly slowed down the spread of the virus and hence the microcephaly epidemic.

To help answer these questions, the Health Secretariat in Rio de Janeiro City provided us with records of all the microcephaly cases and also all the Zika cases up to 26 Sept 2016. Figure 1a and b show the number of microcephaly cases per month from mid-2015 until March 2016 (left) and the number of officially registered cases of Zika per month (right) from 25 October 2015 when it became compulsory to notify the Health Authorities. The key point to note is that babies born with microcephaly from October until the end of January 2016 would have been conceived between December 2014 and March-April 2015, and as Zika is thought to cause most neurological damage in the first 3-4 months [^4^ Based on a recent study in Rio de Janeiro, Brasil et al (2016) report that damage can occur at any time in pregnancy] of the pregnancy (Alvarado & Schwartz, 2016; Cauchez et al, 2016), the officially recorded cases of Zika cannot explain these microcephalic babies. Zika must have been rampant in the city much earlier - as has been suggested by Bastos et al (2016). Our objective is to track the propagation of Zika in the city before its presence had been recognized, by studying the microcephaly cases. So we will focus on microcephaly cases up to the end of January 2016. Another reason for limiting the study to the end of January is that as people became aware that Zika caused microcephaly, women who were thinking of becoming pregnant started modifying their behavior to reduce the chances of catching Zika.

**Fig 1:**
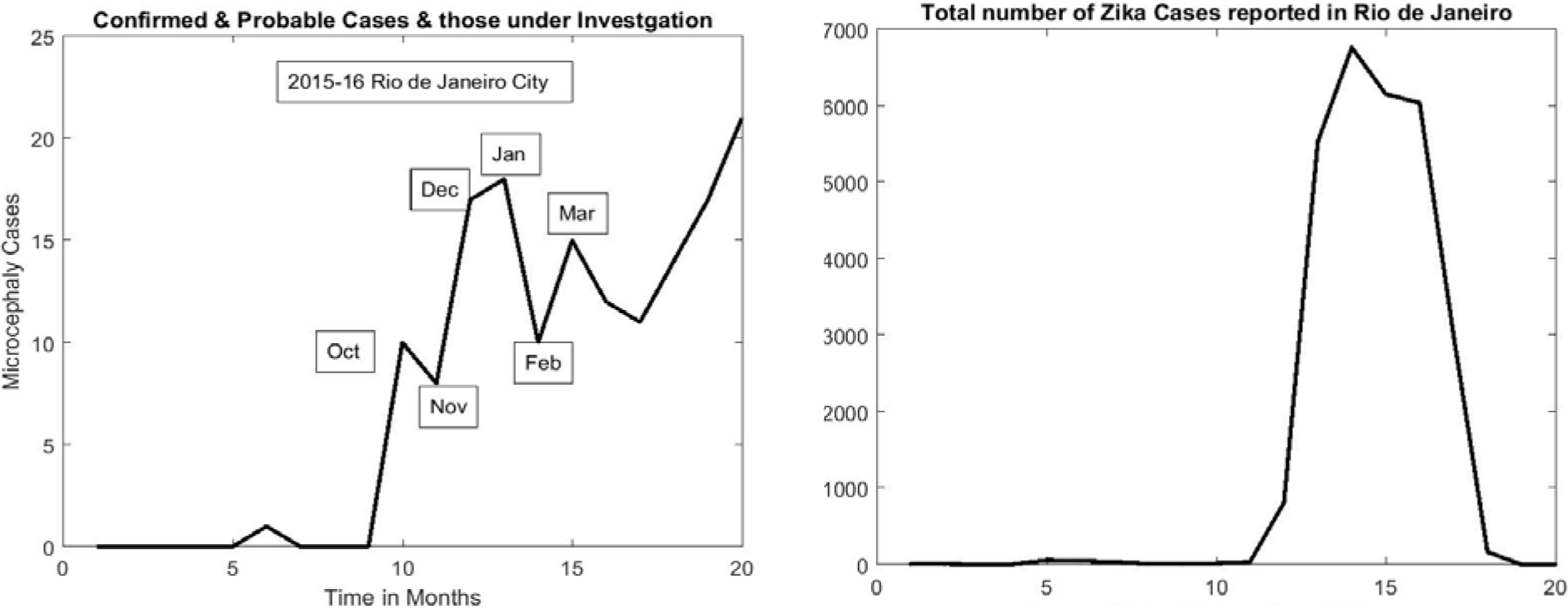
Number of microcephaly cases per month up to March 2016 (left) and the number of registered cases of Zika from October 2015 when it became compulsory to notify the health authorities. As babies born in Oct 2015 were conceived in January and as microcephaly is caused when the mother catches Zika in the first 3-4 months of the pregnancy, the officially recorded Zika cases cannot explain the microcephaly cases. Zika must have been rampant in the city early in 2015.

In the first half of the paper (Section 2) we study how the geographic spread of the microcephaly cases (and hence of Zika) evolved up to the end of January 2016. We show that cases were not evenly spread in proportion to the number of births; they were preferentially located in the northern suburbs apparently following the public transport routes, with virtually no cases in favelas and none in the southern suburbs (Zona Sul). One key difference between the transport systems in the northern and southern suburbs is that the metro & rail system in the north is above ground in the north whereas the metro in the southern part is underground with air-conditioning in carriages and forced ventilation on the platforms. The train system does not extend to Zona Sul.

In the second half of the paper (Section 3) we postulate that the air-conditioning and ventilation prevent mosquitos from biting people who are waiting on platforms in Zona Sul. Agent-based simulations are used to test this hypothesis. After confirming this, we postulate that providing air-conditioning and/or forced ventilation on the rail-metro transport hub in the center of the city (Centro) would significantly delay the propagation of arboviruses in the city, possibly preventing epidemics. The conclusions follow in Section 4.

Our paper is similar to Saad-Roy et al (2016) in that both use the appearance of microcephaly cases as a visible indication of the presence of Zika in a given region. In their paper, the authors establish a mathematical model to describe ZIKV spread from one generic region where there is an epidemic to a region where it has been imported. Their model takes account of both vector transmission and sexual transmission from men to women. Our paper is more specific; we study what actually happened in Rio de Janeiro and we focus on the impact of the transport system on the propagation of the disease. We do not consider sexual transmission.

## Section 2 Analysis of Microcephaly Cases in Rio de Janeiro up to the end of January 2016

Rio de Janeiro is divided into 160 areas called bairros which are regrouped into 33 administrative regions (RA for short). A map showing their layout is given in Fig 2.

**Figure 2:**
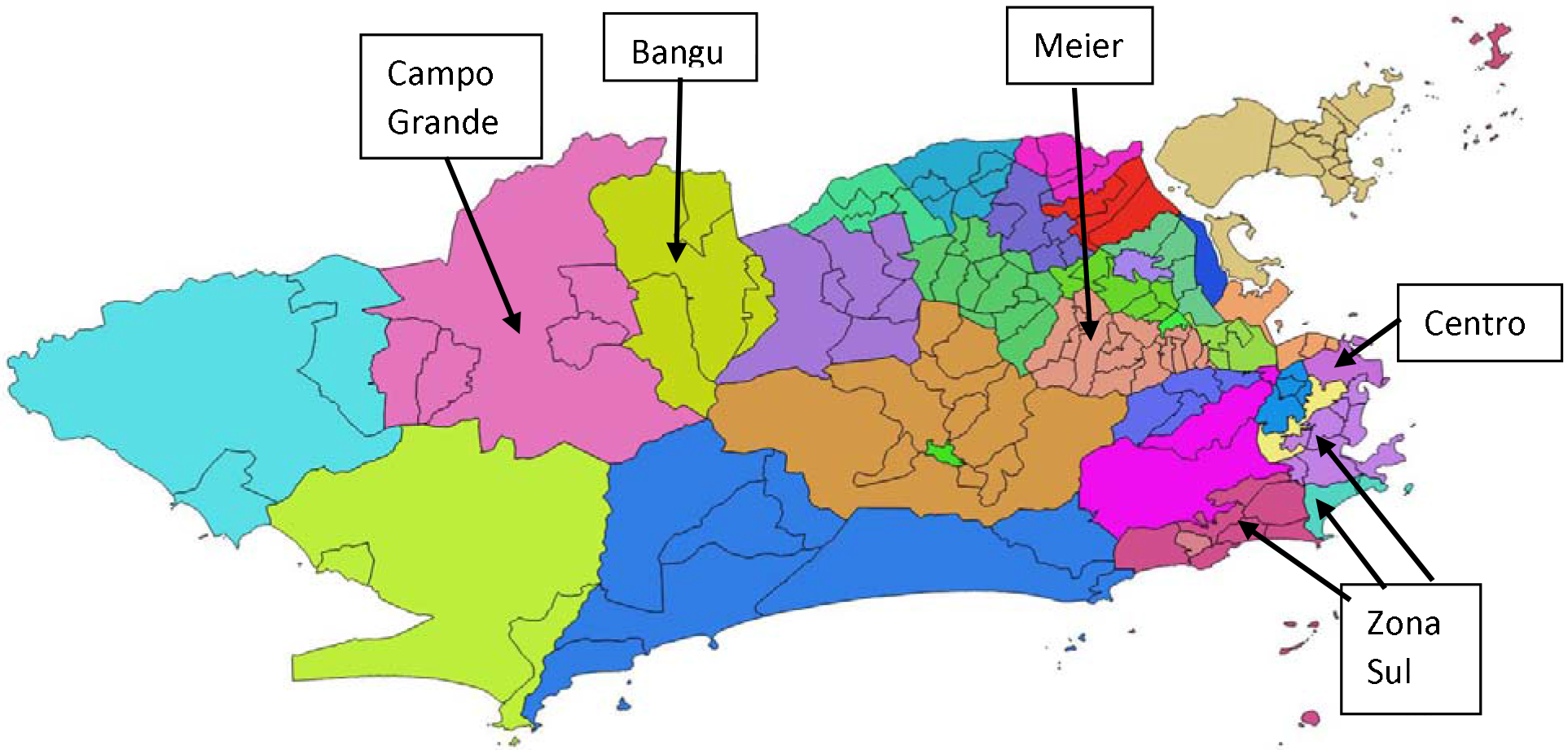
Map showing the 33 administrative regions (RAs) within the city of Rio de Janeiro, with the limits of the bairros inside each one shown as well. The five RAs that will be used for the simulation in the second half of the paper are also shown.

The Health Secretariat of the city of Rio de Janeiro provided us with the data on the 335 cases notified up to 26 Sept 2015. Given the severity of the neurological damage, the Health Department has been carefully monitoring all possible cases in the city. After studying them, some cases were excluded from further study because they do not to fit the criteria for microcephaly, and the remaining 162 cases were divided into three categories: confirmed cases, cases probably due to ZIKV and cases still under study. The confirmed cases were then vetted to check whether the microcephaly could due to another virus. Seventeen of the 84 confirmed cases turned out to be due to other viruses: syphilis (5); herpes (4); toxoplasmosis (4) & cytomegalovirus (4). Table 1 shows the numbers in each class.

**Table 1:**
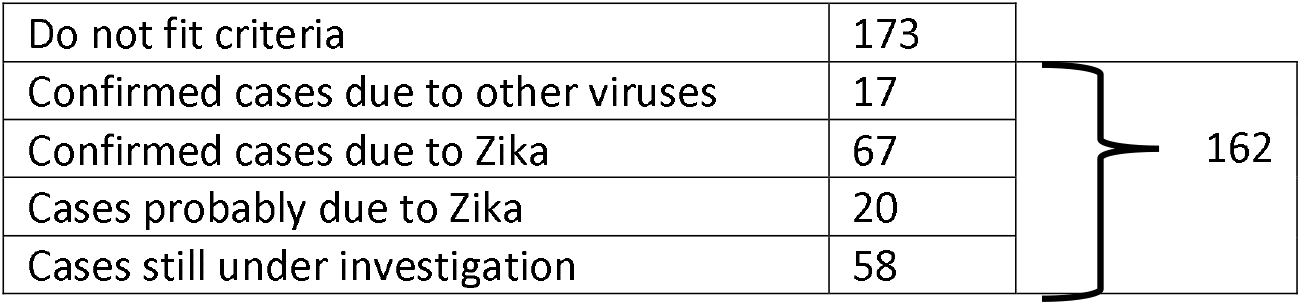
Of the 335 suspected cases of microcephaly that were notified to the health authorities in Rio de Janeiro up to 26 September 2015, only half (162) fitted the criteria, and 17 of these were due to other viruses (syphilis: 5; toxoplasmosis: 4; herpes: 4; cytomegalovirus: 4)

The first step consisted of plotting the date of birth of each microcephaly case as a function of the RA where the baby′s mother lived (Fig 3). Confirmed cases caused by Zika are represented by red circles, probable cases (bright pink), cases still under investigation (yellow), microcephaly cases due to syphilis (bright blue), to toxoplasmosis (green), to herpes (dark blue), to cytomegalovirus (black) and finally those that did not fit the criteria for microcephaly are shown as empty circles. There are few yellow circles corresponding to cases still under investigation up to the end of January; they become much denser from April-May onward. This was one of the reasons for limiting our study up to the end of January 2016. This left us with a total of 47 microcephaly (confirmed, probable and still under study) from Sept 2015 until Jan 2016.

**Figure 3:**
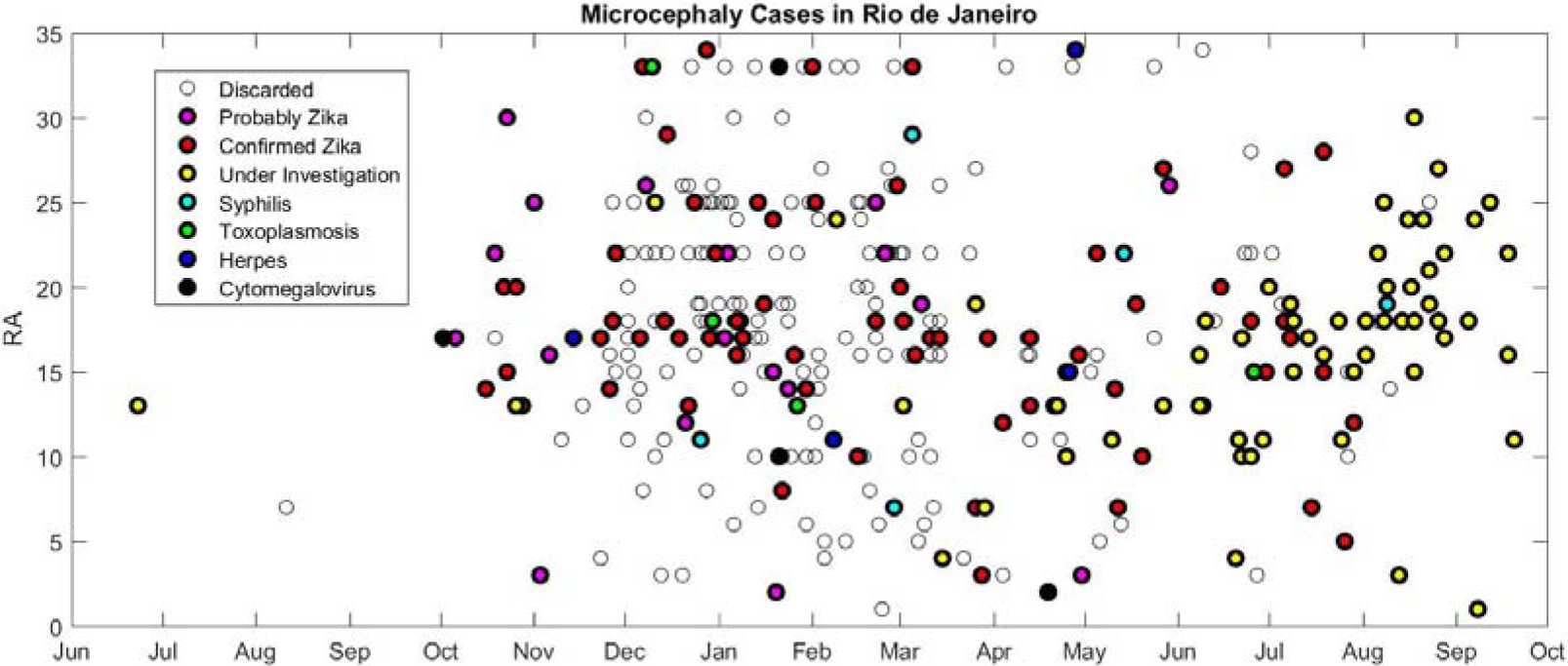
Microcephaly cases in Rio de Janeiro that were notified to the Health Secretariat were analysed and split into 7 categories: confirmed cases caused by Zika (red circles), probable cases (bright pink), cases still under investigation (yellow), microcephaly cases due to syphilis (bright blue), to toxoplasmosis (green), to herpes (dark blue) or to cytomegalovirus (black) and finally those that di not fit the criteria for microcephaly (empty circles). There are few yellow circles corresponding to cases still under investigation up to the end of January; they become much denser from April-May onward.

Figure 4 shows a schematic representation of the layout of the 33 RAs with the number of cases in each up to the end of January 2016. These 47 cases were concentrated in the northern suburbs especially Bangu (7) and Campo Grande (5); there were none in the southern ones called Zona Sul (Lagoa, Copacabana and Botafogo). Next we test whether this spatial distribution could be purely random.

**Figure 4:**
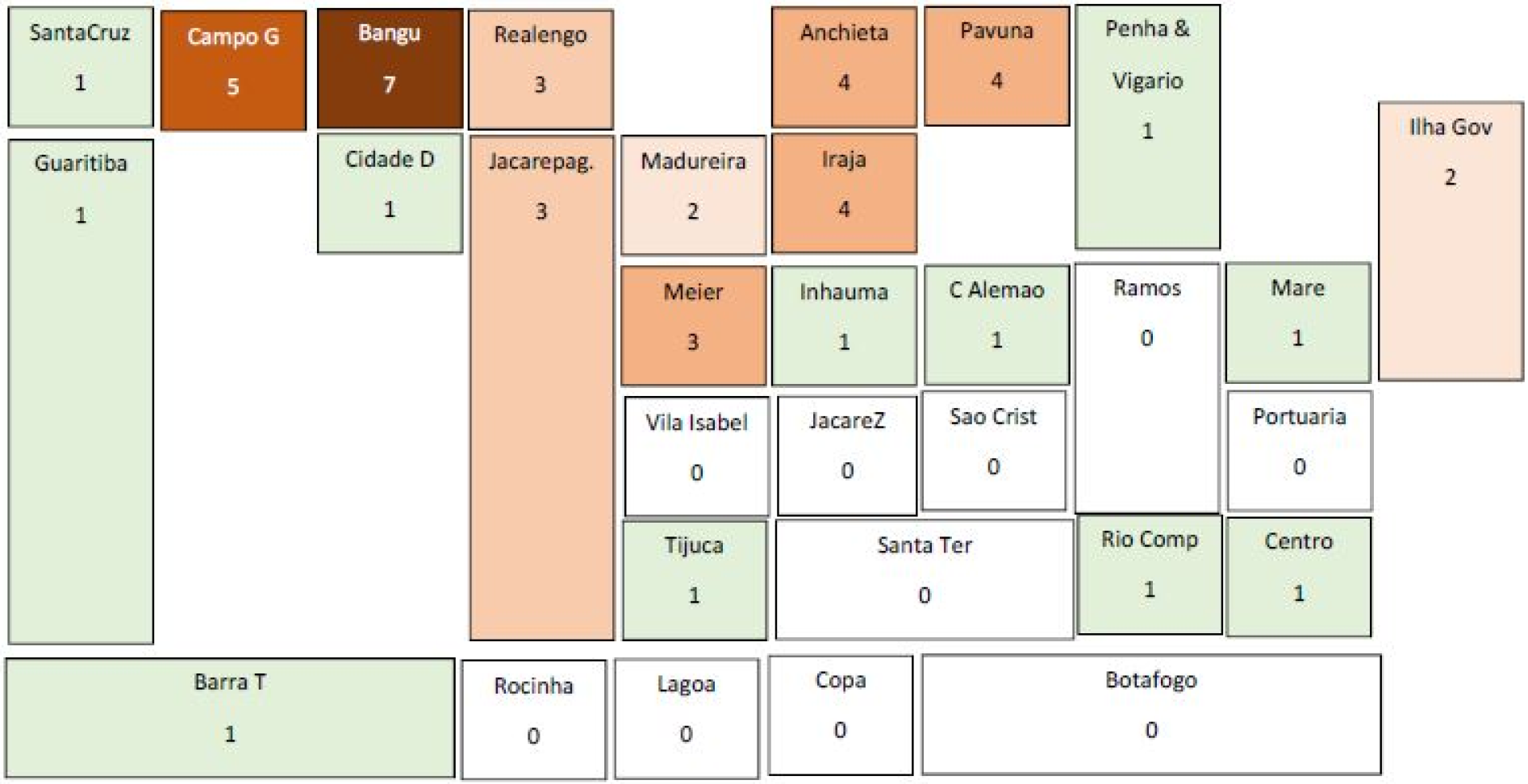
Schematic representation of the layout of the 33 RAs in Rio de Janeiro with the number of cases up to the end of January 2016 (confirmed & probable cases & those still under investigation). Cases are concentrated in the poorer northern suburbs especially Bangu and Campo Grande but there are none in the more affluent Zona Sul (Lagoa, Copacabana and Botafogo). As far as the favelas are concerned, there were no cases in Rocinha and Jacarezinho, and only 1 case in each of the other two important favelas: Complexo da Maré, Complexe do Alemão and Cidade de Deus

### 2.1 Testing whether this spatial distribution could be random

Two separate hypotheses will be tested to determine whether this spatial distribution could have occurred by chance. First we determine the probability of having zero cases in the Zona Sul out of the 47 microcephaly cases if they had been distributed randomly in proportion to the number of live births in the area. This hypothesis is rejected. Next after excluding Zona Sul, we compute the probability of 33 of the 47 cases occurring in the 9 RAs (Santa Cruz, Campo Grande, Bangu, Realengo, Anchieta, Pavuna, Madureira, Iraja & Meier) that lie along the above ground train & metro lines. Again, the hypothesis is rejected.

#### Hypothesis 1: Zero cases of microcephaly could occur at random in the Zona Sul

As there were 948 babies born in Zona Sul from October 2015 to the end January 2016 out of a total of 11812 in the whole city, the probability of 0 cases out of 47 is only 1.96%; so this hypothesis is rejected. See Appendix 1 for details.

#### Hypothesis 2: 33 Cases of microcephaly could occur at random in 9 northern RAs along rail/metro lines

After excluding the 948 babies born in Zona Sul, 10864 had been born in the rest of the city and 5399 were born in the in the 9 RAs (Santa Cruz, Campo Grande, Bangu, Realengo, Anchieta, Pavuna, Madureira, Iraja & Meier) that lie along the above ground train & metro lines. We found that the probability of this occurring at random is only 0.25%. So the hypothesis is rejected too. See Appendix 1 for details.

This shows that this spatial distribution could not occur at random simply in proportion to the number of live births. Clearly some factor (or factors) are causing this. As the northern suburbs are poorer working class areas while the Zona Sul is affluent, the distribution might be controlled by socioeconomic factors.

### 2.3 Testing whether socio-economic factors could explain the spatial distribution

The website of the city of Rio de Janeiro [^5^ http://portalgeo.rio.rj.gov.br/bairroscariocas/index_ra.htm] provides detailed information on the population and the living conditions in each RA, including a Human Development Index (HDI). In the 1990s, the UNDP developed a measure of human development that covered three aspects: longevity, education and economic development (UNDP, 2016; Bitoun, 2010). Here we use the economic index which varies on a scale from 0 to 1 and measures the standard of living. Before starting to test the hypothesis, we plotted the number of microcephaly cases per 1000 live births as a function of the HDI (Fig 5). The yellow diamonds correspond to the 9 northern RAs along the train & metro lines; the red ones are for Zona Sul (2 of the RAs coincide) and the blue ones are for five of the favelas. When we looked at the favelas, we were surprised to find that there were no cases in Rocinha and Jacarezinho, and only 1 case in each of the three densely populated favelas: Complexo da Maré, Complexe do Alemão and Cidade de Deus. The areas with the most microcephaly cases (left column in Table 2) have an intermediate Human Development Index; areas with zero or 1 cases are either affluent areas in the Zona Sul (right column) or favelas which are much more disadvantaged (middle column). Table A3-1 in Appendix 3 lists its value for 31 RA [^6^ The small island llha da Paqueta in the middle of the bay has been excluded. The RAs Penha Viga Geral have been combined as per the city website.] together with the number of children born in the area from October 2015 to January 2016 and the number of microcephaly cases in the same period.

A linear regression model was fitted to test the hypothesis that the number of microcephaly cases per 1000 live births depends linearly on the municipal human development index, against the null hypothesis of a constant. Table 3 shows that this hypothesis is rejected: that is, the number of microcephaly cases per 1000 live births does not depend on the economic human development index.

**Table 2:** The Municipal Human Development Index^3^ in selected RAs. The areas with most microcephaly cases (left column) are poorer working class areas: not affluent areas (right) nor favelas (middle column)

| RA (north) | Human Dev Index | RA favelas | Human Dev Index | RA Zona Sul | Human Dev Index |
| --- | --- | --- | --- | --- | --- |
| Bangu (7) | 0.72 | Rocinha (0) | 0.67 | Botafogo (0) | 0.99 |
| C. Grande (5) | 0.73 | Jacarezinho (0) | 0.64 | Copacabana (0) | 1.00 |
| Realengo (3) | 0.75 | C. Alemão (1) | 0.64 | Lagoa (0) | 1.00 |
| Anchieta (4) | 0.72 | C. Mare (1) | 0.65 |  |  |
| Pavuna (4) | 0.69 | Cidade Deus (1) | 0.66 |  |  |

<sup>5</sup> http://portalgeo.rio.rj.gov.br/bairroscariocas/index_ra.htm
<sup>6</sup> The small island Ilha da Paqueta in the middle of the bay has been excluded. The RAs Penha Viga Geral have been combined as per the city website.

**Figure 5:**
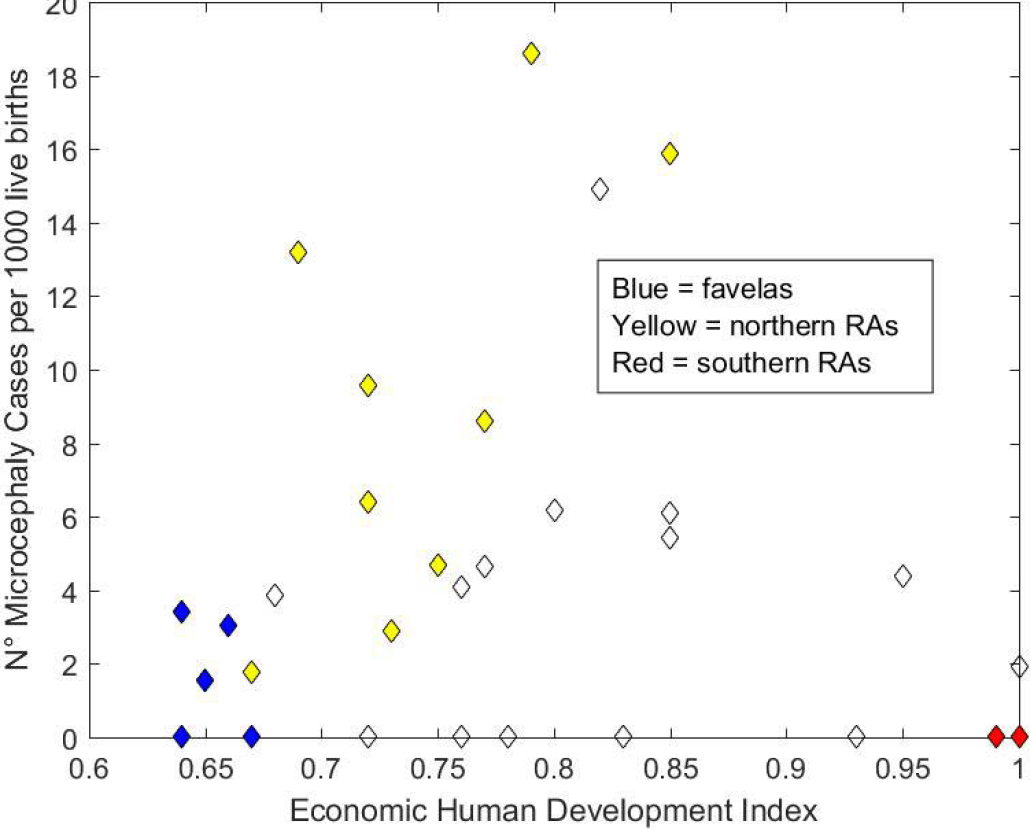
The number of microcephaly cases per 1000 births in each of the RAs in Rio de Janeiro, plotted as a function of the Municipal Human Development Index. The 5 dark blue diamonds correspond to favelas; the 9 yellow diamonds represent the RAs along the northern train & metro lines while the 2 red ones correspond to the three RAs in Zona Sul (2 of these coincide).

**Table 3:**
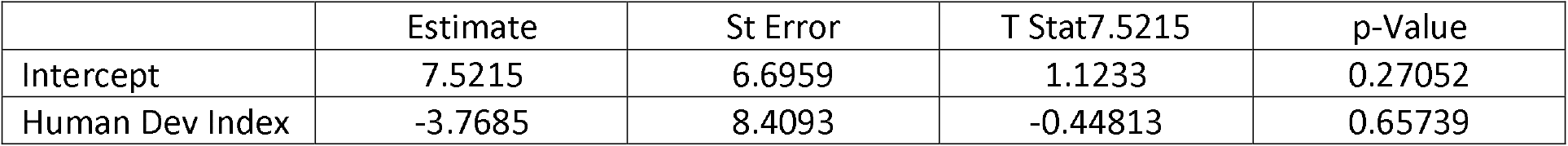
Fitted Linear Regression Model of the Number of Microcephaly cases per 1000 Live Births as a Function of the Municipal Human Development Index

### 2.3 Rail & metro system in Rio de Janeiro

The public transport in Rio de Janeiro consists of train lines which are above ground and which take passengers from the northern suburbs into the city center (Fig 6) and metro lines which are above ground from the city center out north to Pavuna and below ground from the city center to Zona Sul in the southern suburbs (Fig 7), together with bus routes.

Anyone who travels on the metro in Rio notices how cold it is in the carriages and how windy it is on the platforms. We were curious to find out why it had been built this way. Was the aim to keep mosquitos away or was this just a lucky consequence? The answer was provided by a former civil engineer who had worked on designing the stations soon after graduating. It had nothing to do with mosquito prevention. The fans and the air-conditioning in the carriages are for passengers’ comfort. Added to this, when trains go through tunnels, they push a wedge of air through in front of them, rather like a piston does, and large fans on the platforms increase the circulation of the air. A side effect is that it makes it difficult for mosquitos to bite people.

In the next section 3 we propose to use agent-based simulations to test whether eliminating mosquitos from the metro stations in Zona Sul (the southern suburbs) would slow down the propagation of Zika and hence could explain why there are so few microcephaly cases in that part of the city.

**Figure 6:**
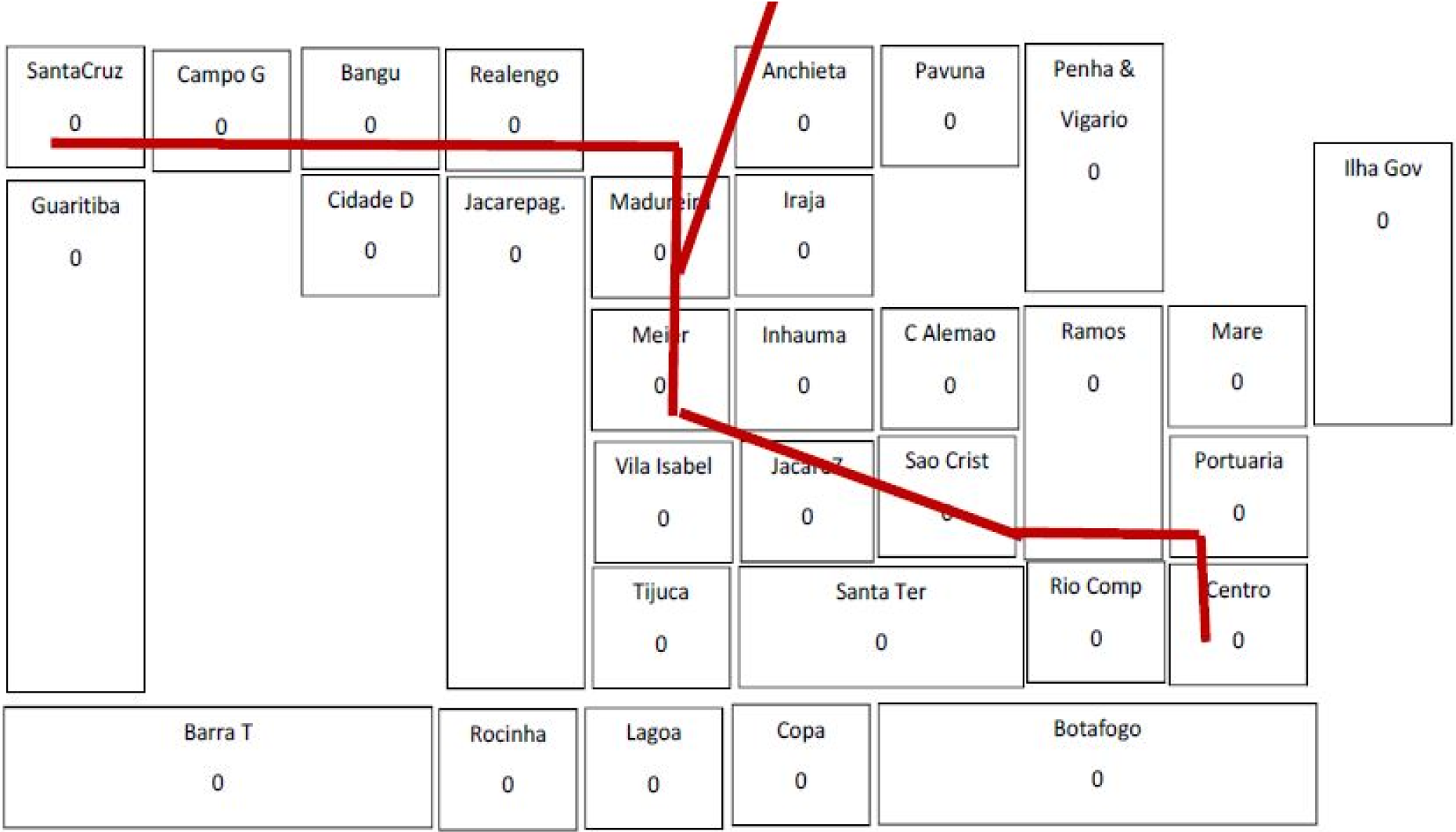
Schematic layout of the RAs showing the train lines which are above ground and which take passengers from the northern suburbs into the central station. No train lines go to the southern suburbs.

**Figure 7:**
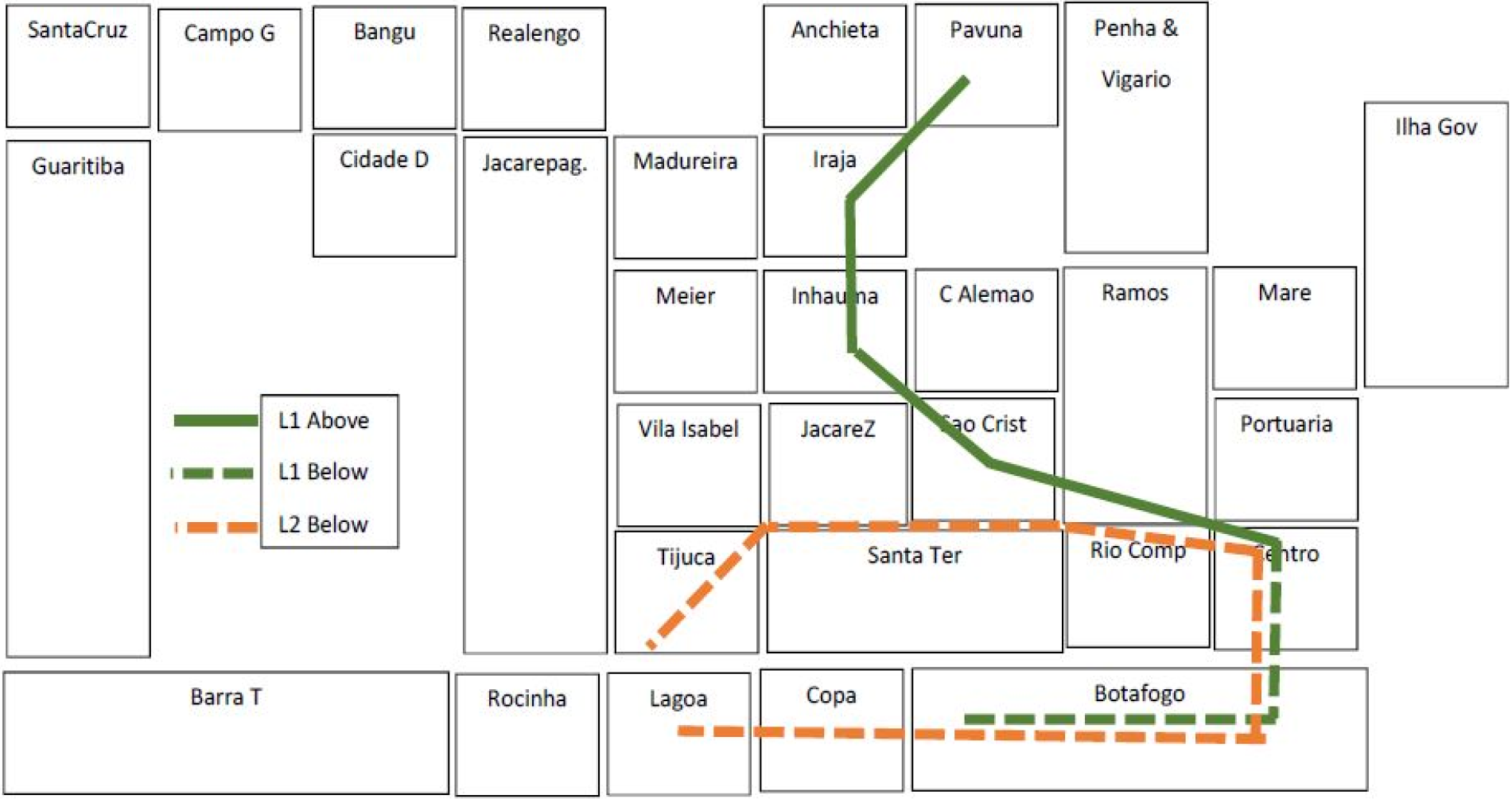
Schematic layout of the RAs showing the metro lines. The solid lines correspond to ones that are above ground, while the dotted ones are underground. The platforms in the underground stations have forced ventilation and the carriages are air-conditioned.

## Section 3 Simulating the Impact of the Transport System on the Propagation of Zika

It is widely recognized that human mobility is an important factor in spreading arboviruses such as dengue fever, chikungunya and zika (Caraco et al, 2001; Barmak et al, 2011; Barmak et al, 2016).

We want to test the hypothesis that air-conditioning in metro carriages and ventilation on the metro platforms in Zona Sul has made it more difficult for mosquitos to bite people waiting on the platforms and hence led to a decrease in microcephaly cases in the southern suburbs. We propose to use agent-based simulations to test this on a simplified (“toy”) example consisting of 5 RAs in Rio de Janeiro:

- three in the northern suburbs along the above ground train lines (Bangu, Campo Grande & Meier),
- the city center with the main business area (Centro)
- and the southern suburbs (Zona Sul).

Following Barmak et al (2011) and Barmak, Dorso & Otero (2016) we divided each of these 5 RAs into two areas: the area around the train/metro station and the rest of the RA. This gives as 10 sub-zones each with its own mosquito population. There is some debate as to how far Aedes Aegypti mosquitos fly. Some authors believe that they only fly short distances from their emergence site (Reuben et al, 1975; Trpis & Hausermann, 1986; Service 1997) but Honorio et al (2003) found them up to 800m from the release point. For simplicity, we assume the 10 mosquito populations are distinct and do not mix. Table 4 gives the numbers of inhabitants in these five areas at the time of the 2010 census. As very few people actually live in the city center in Rio de Janeiro compared to the other four areas, we ignore their presence in our model.

**Table 4:** Population in each of the areas in the “toy” example, from the 2010 census [^7^ http://portalgeo.rio.rj.gov.br/bairroscariocas/index_ra.htm]

| RA | Bangu | C. Grande | Meier | Centro | Botafogo | Copacabana | Lagoa |
| --- | --- | --- | --- | --- | --- | --- | --- |
| Pop | 413,000 | 542,000 | 398,000 | 41,000 | 240,000 | 161,000 | 168,000 |
|  |  |  |  |  | Zona Sul = 569,000 |  |  |

Following Barmak et al (2011) and Barmak, Dorso & Otero (2016), the populations living in the four remaining areas are split into two groups: those that go to the city center to work each day and those who remain in the home area. The latter go to work or to school there or they stay at home. People who stay in their residential area mix freely with each other but not with those from other areas. In contrast, people who go to work in the business area, mix with others there while doing business and while having lunch, and then again on the platform while waiting to go home.

### 3.1 Interactions between mosquitos & people

In contrast to Barmak and colleagues who only consider one time period per day, we divide each day into 4 periods: early morning (when those who go to the city center wait on the platforms & get bitten by the station mosquitos), midday (when people are in contact with either those in the city center, or with those in their home area), late afternoon when the travellers wait on the platforms at Centro & are bitten by that population of mosquitos) and finally the evening when everyone is at home & gets bitten by the population of mosquitos in that area. Table 5 summarises the interactions between the 10 mosquito populations (M1 to M10) and the 8 populations of people (P1 to P6, P9 & P10). For example, the populations P1 + P3 + P5 + P9 wait for a train home late in the afternoon can be bitten by mosquitos from the M7 population. In the evening, populations P1 + P2 are at home in the Bangu area & can be bitten by mosquitos from population M2.

**Table 5:** Summary of the interactions between the 10 mosquito populations (M1 to M10) and the 8 groups of people (P1 to P8) at the four different periods in the day.

|  | Bangu Station<br>M1 | CampoGrande<br>Station M3 | Meier Station<br>M5 | Centro Station<br>M7 | Zona Sul Area<br>M9 |
| --- | --- | --- | --- | --- | --- |
| Morning | P1 | P3 | P5 |  | P9 |
| Midday |  |  |  |  |  |
| Afternoon |  |  |  | P1+P3+P5+P9 |  |
| Evening |  |  |  |  |  |
|  | Bangu Area<br>M2 | Campo Grande<br>Area M4 | Meier Area<br>M6 | Centro Business<br>Area M8 | Zona Sul Area<br>M10 |
| Morning | P2 | P4 | P6 |  | P10 |
| Midday | P2 | P4 | P6 | N°1+N°3+N°5+N°9 | P10 |
| Afternoon | P2 | P4 | P6 |  | P10 |
| Evening | P1+P2 | P3+P4 | P5+P6 |  | P9+P10 |

To start the simulation we inject two infectious people into Bangu (because it had the most cases of microcephaly); one person remains within the area during the day and the other goes to work in the city center. An infectious person on the Bangu station could transmit Zika to the M1 mosquito population in the morning, then to the M8 population in the business area in the middle of the day, then to the M7 population at the Centro station and finally to the M2 population in the Bangu residential area. After the incubation period, the M7 & M8 mosquitos could transmit to virus to people from other areas (P1, P3, P5 & P9) and so the disease is propagated.

### 3.2 Parameters in the model

We consider a SEIR model (susceptible, exposed, infectious and recovered) for people and a SEI model for mosquitos. Mosquitos do not recover; but they do die. As we only consider a period of several months, we ignore human mortality and people moving their residence from one area to another. Moreover, when a susceptible mosquito bites an infectious person the virus is not always transmitted to it, and similarly when an infectious mosquito bites a susceptible person. These transmissions are controlled by two probabilities. The main parameters in the model as listed in Table 6 were taken from Bastos et al (2016). When a mosquito dies, it is replaced by a new susceptible mosquito. We do not allow for vertical transmission between mosquitos and their progeny. For simplicity, the incubation periods and the time to recovery are taken to be deterministic.

**Table 6:** The main parameters in the model together with their values

|  |  |  |
| --- | --- | --- |
| Probability human to mosquito | Probability mosquito to human | Daily mortality rate mosquitos |
| 0.2 | 0.2 | 0.05 |
| Incubation period in humans | Time before recovery in humans | Incubation period in mosquitos |
| 5 days | 6 days | 7 days |

### 3.3 Simulation procedure

In our toy example we assume that there are 10 mosquitos in each of the 10 populations M1 to M10 and that there are 10 people in the each populations P1 to P6, P9 and P10. So there are 20 inhabitants in Bangu in the model: 10 who travel to the city center each work day and 10 who remaining in the Bangu area. As we have chosen to ignore people living in the city center, the populations P7 & P8 are zero.

The flow-chart for the multi-agent simulations is given in Appendix 2. It was performed for a total of 120 days, firstly with 10 mosquitos in each of the 10 populations and then again with a zero population on the platforms of the metro stations in Zona Sul. To ensure that the results can be compared meaningfully we used exactly the same values for the three random variables: the mosquito mortality, the success/failure of the virus transmission from an infectious human to a susceptible mosquito, and vice versa.

### 3.4 Impact of removing the mosquitos from the Zona Sul platform

To illustrate the impact of removing the mosquito population (M9) from the platform at the Zona Sul metro station while holding all the other parameters fixed and using **exactly the same random variables**, Fig 8 shows the evolution of (1) the number of recovered cases of a total of 160 as a function of time. The red line corresponds to the base case where there are mosquitos on the Zona Sul platform, the black one to the case without mosquitos. The impact of removing mosquitos is quite clear, but small overall. Next we compare the number of recovered cases, area by area, for Zonal Sul (Fig 9), for Bangu and Campo Grande (Fig 10a & b). In this case removing the mosquitos lead to a dramatic drop in the number of cases in Zona Sul: 50%-60% less from Day 60 to Day 90, but it produced no changes in the other areas; the black line coincided with the red one which is no longer visible.

Having seen how effective this strategy is overall, we were curious to see what happened in Zona Sul. Fig 11 shows that eliminating the mosquitos on the Zona Sul platform is more effective in the short term from Day 60 to Day 90 but less effective afterwards.

**Figure 8:**
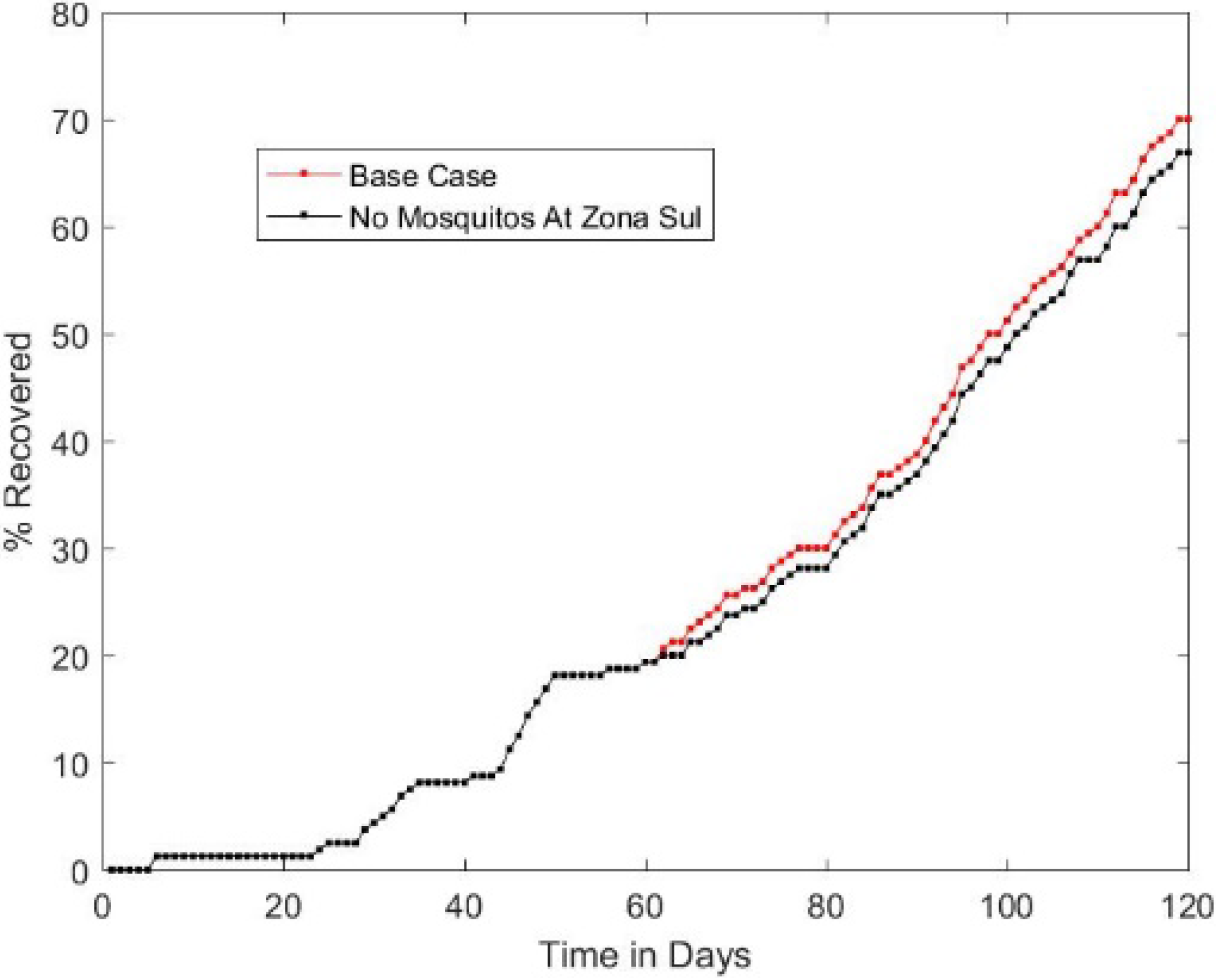
The evolution of the number of recovered cases (out of a total of 160) with mosquitos on the Zona Sul platform (red) and without (black) as a function of time after the arrival of 2 infectious people in Bangu: one who stays in the Bangu area and one who travels to the city center each work day.

**Figure 9:**
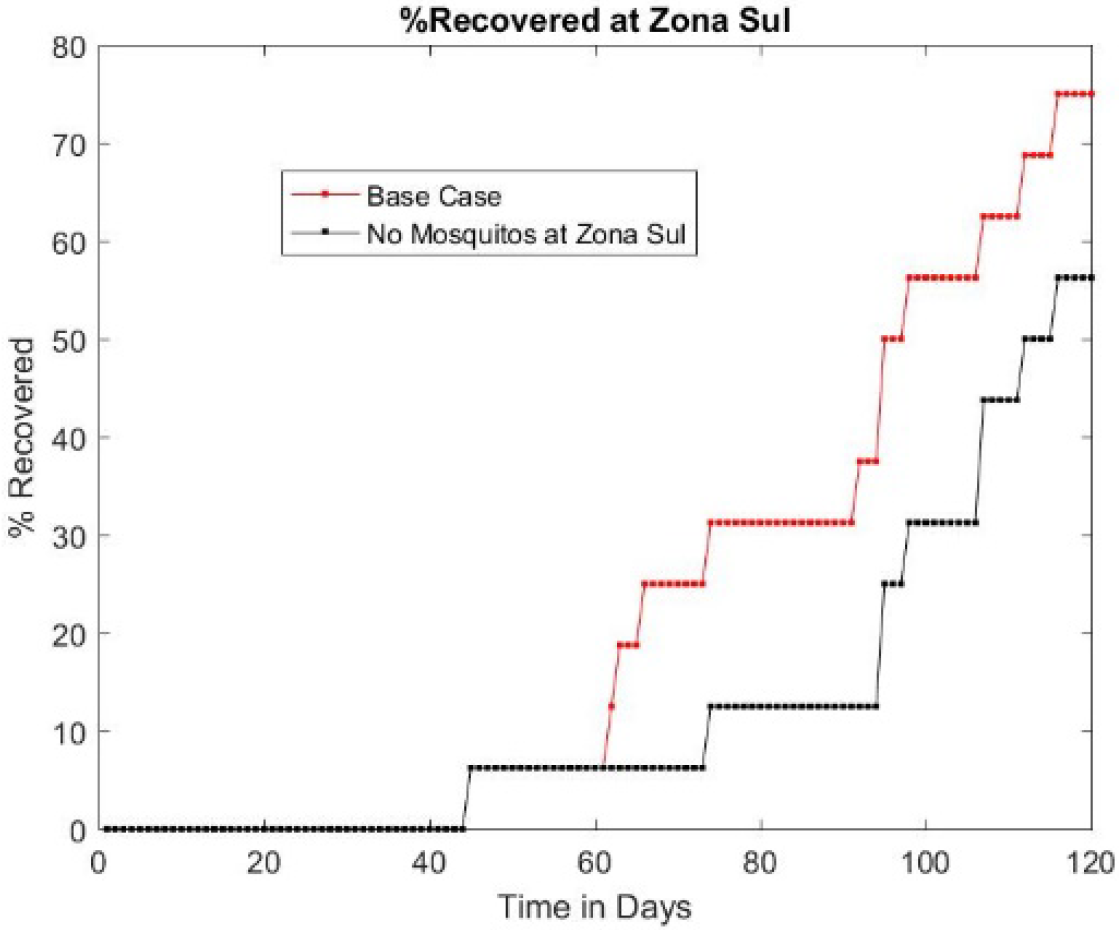
The evolution of the number of recovered cases in Zona Sul (out of a total of 40) as a function of time. The redline corresponds to the case where there are mosquitos on the Zona Sul platform, the black one to the case without mosquitos. The impact of removing mosquitos is quite clear. In this case it halved the number of cases from day 60 to day 90 when the epidemic would be taking off otherwise.

**Figure 10a & b:**
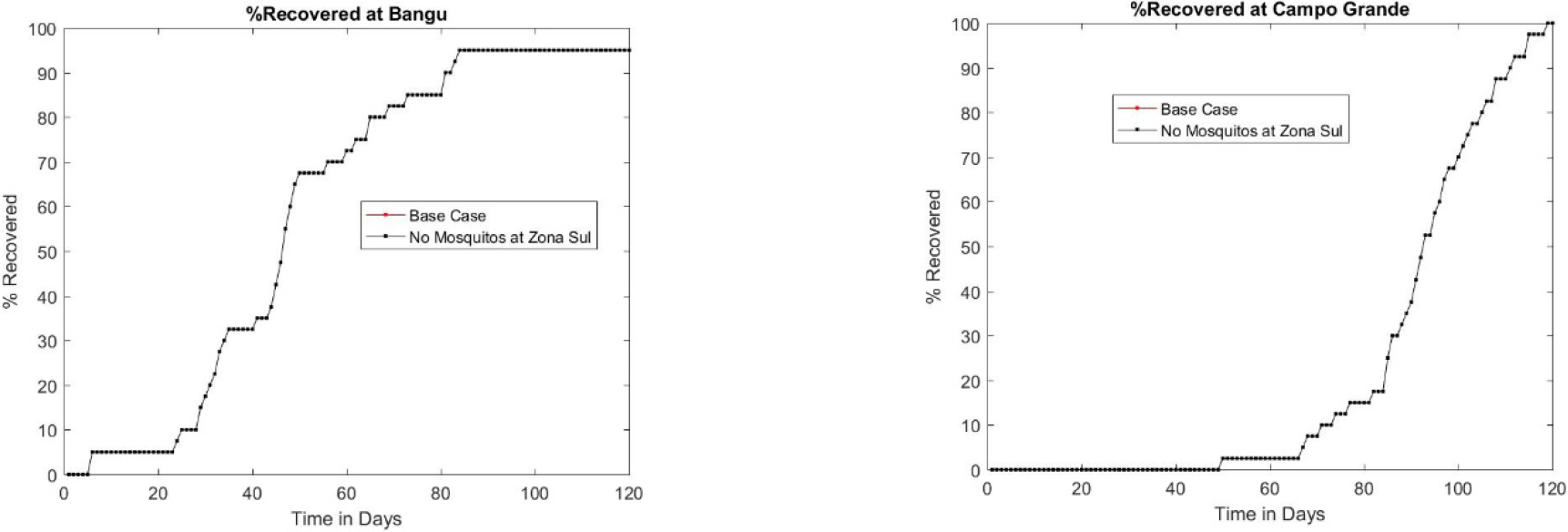
The evolution of the number of recovered cases in Bangu (left) and Campo Grande (right) as a function of time. The red line coincides with the black one. Removing mosquitos at Zona Sul did not affect the evolution of the epidemic in other areas.

### 3.5 Impact of removing the mosquitos from the platform at Centro

Having seen the impact of removing the mosquitos from the Zona Sul platform, we postulate that removing them from the city centre rail hub, Centro, might be even more effective in slowing down the progression of the epidemic. Fig 11 shows that this is the case: the percentage of recovered cases after 120 days is about 50% compared to 68% – 70% before. So removing the mosquitos from the city center rail hub, Centro, has led to a 25% drop in cases. The mosquitos can be removed by putting fans on the platforms or by covering the area & installing air-conditioning. Alternatively insecticides could be used. The first option is more friendly from an ecological point of view.

**Figure 11:**
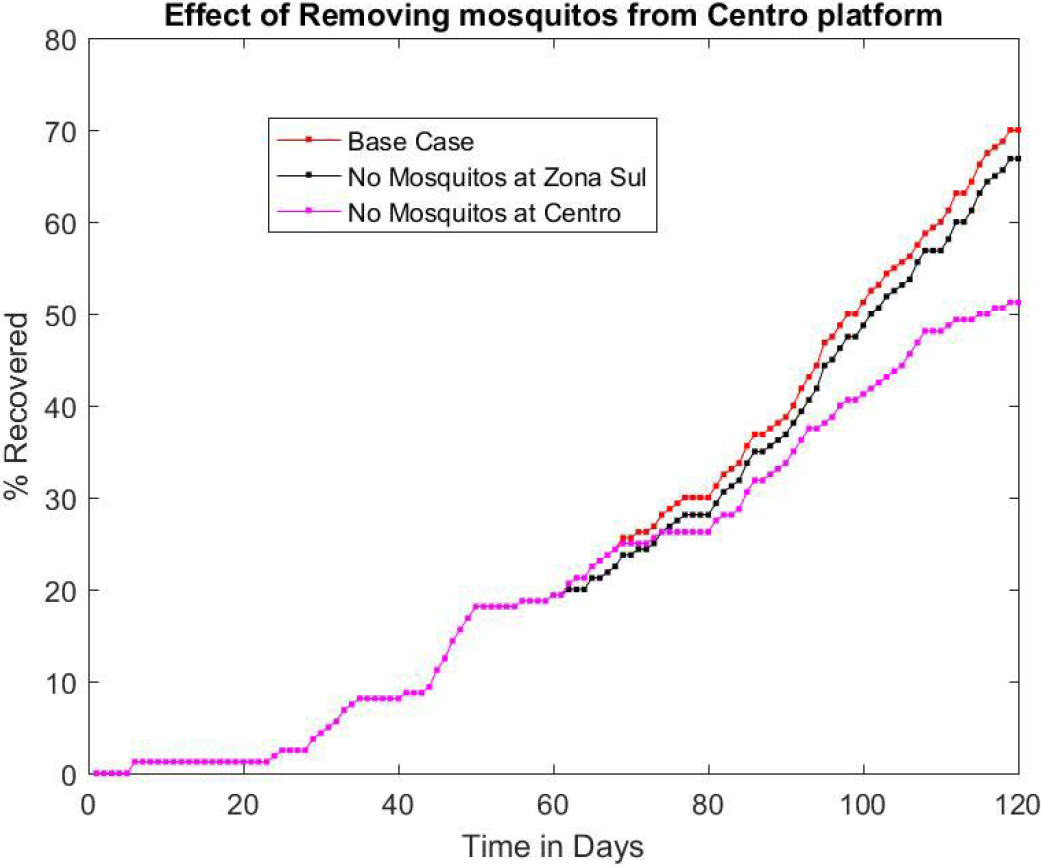
The evolution of the number of recovered cases (out of a total of 160) as a function of time. The redline corresponds to the case where there are mosquitos on all the platforms, the black one to the case with no mosquitos on the Zona Sul platform and the mauve line to the case with no mosquitos on the platform at Centro. The impact of removing the mosquitos from the rail hub is quite clear. It leads to a drop in the number of cases from day 60 onwards, going up to 25% by day 120.

**Figure 12:**
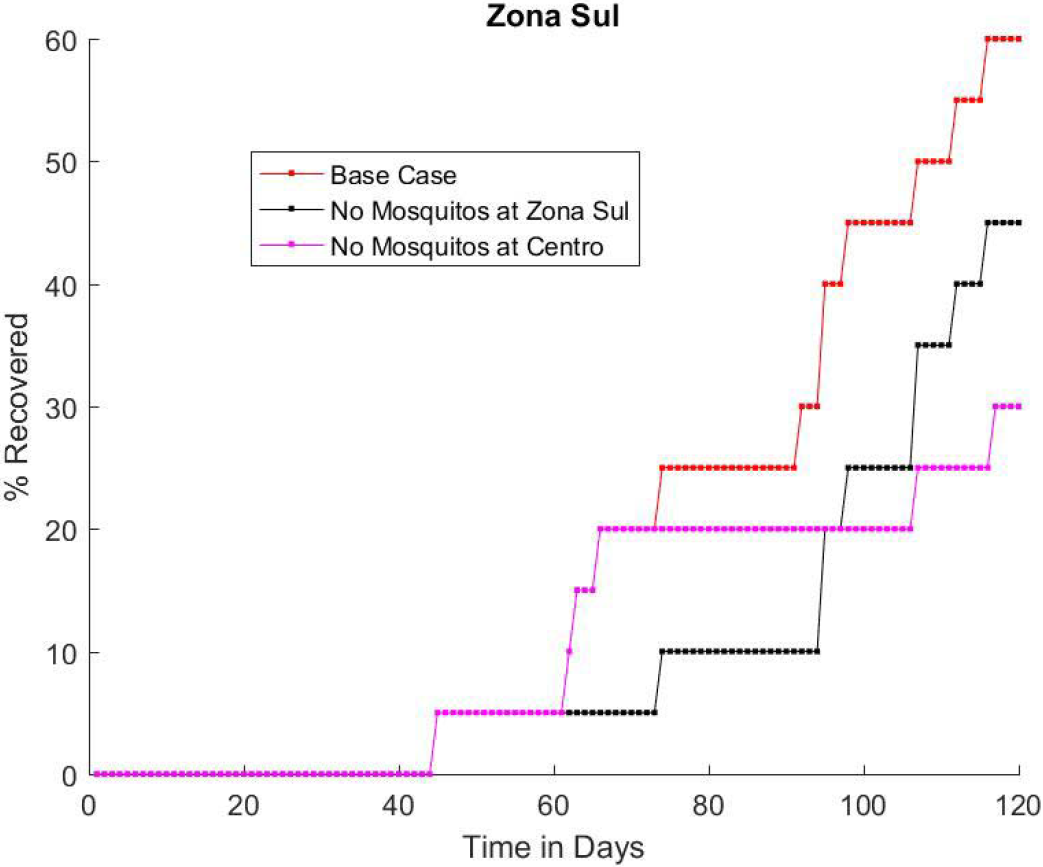
The evolution of the percentage of recovered cases in Zona Sul as a function of time. The red line corresponds to the case where there are mosquitos on all the platforms, the black one to the case with no mosquitos on the Zona Sul platform and the mauve line to the case with no mosquitos on the platform at Centro. Removing the mosquitos from the rail hub leads to a drop in the number of cases in Zona Sul compared to the Base Case from Day 70 onwards, and compared to the second case from Day 95 onward. In the short term it is more efficient to eliminate the mosquitos from the platform at Zona Sul itself, but not in the long term.

### 3.6 Box Plots for the Base Case and the Two Alternatives

Although Figures 8 - 12 illustrate the impact of removing the mosquitos from the platforms at Zona Sul or at Centro, they were obtained from a single set of simulations. Next we need to see what happens for a reasonably large number (100) of sets of simulations. First we computed the median number of recovered cases after 120 days (October 2015 to end January 2016), for the base case and the one with no mosquitos. This confirms what was seen in the simulation used as an illustration (Fig 9).

**Table 7:** Median number of recovered cases after 120 days (equivalent to Oct 2015 – Jan 2016)

|  | Base Case | No Mosquitos on Zona Sul Platform |
| --- | --- | --- |
| Median | 7.5 cases (18.75%) | 13 cases (32.5%) |

Fig 13 shows the box plots for the total percentage of recovered cases for the base case and the two alternatives, with the base case (left), the case with no mosquitos on the Zona Sul platform (middle) and the one with no mosquitos on the Centro platform (right). These confirm that removing the mosquitos from the platform at Centro leads to a more significant decrease in the total number of cases. Note that in all three cases, there were simulations where there were only 2 recovered cases. That is, no cases other than the additional index cases occurred.

**Figure 13:**
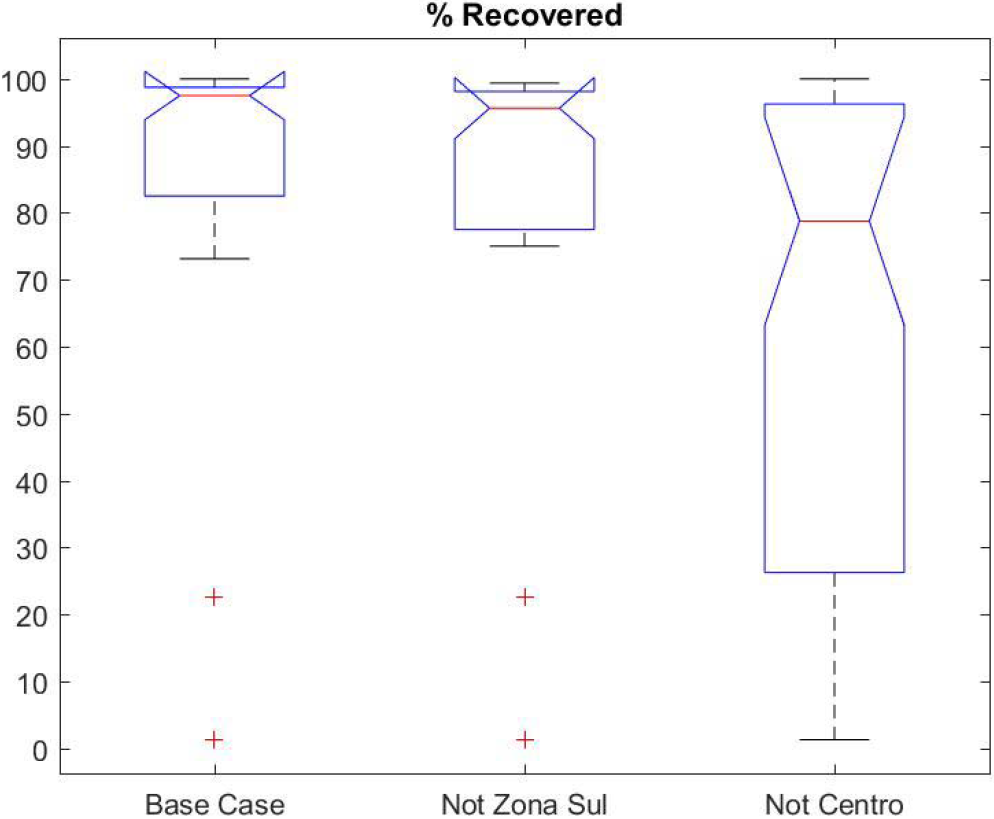
Box plots of the total percentage of recovered cases for the three cases, base case (left), case with no mosquitos on the Zona Sul platform (middle) and no mosquitos on the Centro platform (right). Removing the mosquitos from the platform at Centro leads to a more significant decrease in the total number of cases. Note that in all three sets of simulations, there were simulations where there were only 2 recovered cases. These correspond to the index cases for the simulation. This means that the virus was not transmitted to any other people.

## 4. Discussion & Conclusions

From October 2015 to the end of January 2016, 47 cases of microcephaly that were not due to other viral infections were observed in the city of Rio de Janeiro. These children were conceived from Dec 2014 to April 2015, far too early to be explained by the officially recorded cases from October 2015 onward. Zika must have been rampant in the city from late 2014 onward. The cases could have been asymptomatic or mild enough so that the person did not seek treatment, or if they saw a doctor, the case could have been mistaken for dengue fever.

Most of the 47 microcephaly cases were preferentially located in the northern suburbs, with no cases in the southern ones (Zona Sul) and very few in the favelas. After testing the hypothesis that this spatial distribution was due to chance and having rejected it, we next tested whether it was due to socioeconomic factors: Zona Sul is an affluent area with a high standard of living (human development index) whereas the northern suburbs are working class. But as can be seen from Fig 5, the favelas have an even lower standard of living and yet there were very few cases there. So not surprisingly this hypothesis was also rejected.

After looking at the layout of the public transport system (Figs 6 & 7) and knowing that the metro lines in the southern part of the city are underground whereas the rail and metro lines in the northern part are above ground, we postulated that the absence of cases in Zona Sul and the concentration in the northern suburbs along the train & metro lines was due to the public transport system. The underground metro lines have strong ventilation on platforms and chilly air-conditioning in the carriages, making it difficult for mosquitos. In contrast, people who are waiting for trains above ground can easily be bitten by mosquitos.

The question is: Would the absence of mosquitos on metro platforms in the Zona Sul area be enough to slow down the arrival of Zika by several months? To test this idea, we set up an agent-based simulation model of a small but representative part of the city. It consists of three northern suburbs (Bangu, Campo Grande and Meier) which lie along the train line to the business area in the city center (Centro) plus the Zona Sul. As the populations in the three northern suburbs and in the Zona Sul are between 400,000 and 550,000 compared to only 40,000 in Centro we ignored the city center population. In each of the 4 areas we divided the inhabitants into two groups: those who go to the city center for work each day and those who remain in their home suburb (going to work or school, or staying at home). This gives 8 human populations. As mosquitos do not fly far compared to the distance people go by public transport, we split the mosquitos in each suburb into those on & around the station and those in the rest of the suburb and assumed that the two groups do not mix.

In our model, the day is split into 4 time periods (morning, midday, afternoon & evening). In each period mosquitos select a person at random to bite: if the mosquito is infectious the person may be exposed to the virus and conversely if the mosquito was susceptible and the person was infectious, the mosquito may become exposed. So our model is a stochastic SEIR model for the humans but a SEI model for mosquitos. Mosquitos die at random but do not recover.

The agent-based model shows that having no mosquitos biting people on the Zona Sul platform halves the number of Zika cases in Zona Sul in the first 90 days after the index case was introduced in the northern suburb of Bangu (Fig 9). As expected, no effects were observed in the other areas. The median number of cases after 120 days (i.e. 4 months) was also computed based on 100 simulations. We therefore conclude that the strong ventilation on the underground metro platforms together with the air-conditioning in carriages could explain the lack of microcephaly cases in Zona Sul up to the end of January. (One or two occurred in subsequent months). This is a very encouraging result. It suggests that removing mosquitos from transport hubs would lead to an even more dramatic reduction in the propagation of arboviruses.

So next, we tested what would happen in our model if the mosquitos were removed from the platform at the transport hub in the city center would have. As expected, the impact turns out to be much more marked than just for Zona Sul (Fig 12 for the reference simulation; Fig 13 on average over 100 simulations).

### Recommendation

We therefore recommend that strong ventilation be installed at the rail/metro hub at Centro and also at the nearby bus hub (Rodoviaria). Judging from our agent-based simulations this could produce a very positive effect, without the disadvantages of using insecticides.

Although the results are specific to the city of Rio de Janeiro, the method used could be used to model the propagation of vector-borne epidemics in large cities where public transport is important, for example, cities in southern & central America, in Asia, in Africa and around the Mediterranean basin. One of the method′s advantages is that the hypothesis of perfect mixing is not required. The metropolitan area can be split into suburbs where the mixing hypothesis is more reasonable. Moreover the impact of the transport system can be modeled specifically.

### Limitations of this study

The results presented were obtained from a simplified model of human mobility in Rio de Janeiro.

Several aspects in the agent-based simulations could easily be improved in the future.

1. We ignored vertical transmission of virus from female mosquitos to their progeny.
2. We ignored the mobility of the mosquitos themselves.
3. We limited the simulation study to 5 parts of the city (RAs). In the future this could be extended to cover the whole city.
4. Most important of all we only considered trips between the city center and the 4 suburbs (Bangu, Campo Grande, Meier and Zona Sul). No travel between adjoining suburbs was considered. It would not be difficult to take account of trips between suburbs as well as those to the city center.

## Appendix 1 Testing two hypotheses about random distribution of microcephaly cases

Here we test two hypotheses about this spatial distribution

1. What is the probability of having zero cases in the Zona Sul out of the 47 microcephaly cases if they had been distributed at random in proportion to the number of live births in each RA.

  Pr (Born in Zona Sul) = 948/11812 = 0.0803
  Pr (0 cases in Zona Sul) = (0.9197)^47^= 0.0196 1.96%< 5% Consequently we reject the hypothesis that 0 cases in Zona Sul could arise at random.
2. Now we consider the 9 RAs (Santa Cruz, Campo Grande, Bangu, Realengo, Anchieta, Pavuna, Madureira, Iraja & Meier) in the northern suburbs where the train & metro lines are above ground. Thirty-three out of the 47 microcephaly cases occurred in these RAs. What is the probability after excluding the births in Zona Sul, that 33 of the 47 cases could have occurred in these 9 RAs?

|  |  |  |
| --- | --- | --- |
|  | City of Rio de Janeiro | Zona Sul |
| N° of Babies Born Oct 15 – Jan 16 | 11812 | 948 |

  Pr (Born in 9 RAs) = 5399/10864 = 0.497 As for the first hypothesis the number of cases is a binomial random variable. We will use a normal approximation to the binomial distribution to compute the probability of 33 cases or more out of 47.
  Mean µ =47 ⨯ 0.497 = 23.36
  Standard deviation σ = √(47⨯0.497⨯0.503) =3.43
  Pr (33 out of 47 cases) = N((33-23.36)/3.43) = N(2.81) = 0.0025 Again, the hypothesis is rejected as the probability is only 0.25%.

|  |  |  |
| --- | --- | --- |
|  | City excluding Zona Sul | 9 RAs listed above |
| N° of Babies Born Oct 15 – Jan 16 | 10864 | 5399 |

**Appendix 2.** The number of cases of microcephaly per RA from Oct 2015 to January 2016, the Municipal Human Development Index for each RA, the number of live births for the same period and the number of cases per 1000 live births.

| Name | N° Cases | Human Dev Index | N° Births | Cases per 1000 Births |
| --- | --- | --- | --- | --- |
| Porturia | 0 | 0.72 | 144 | 0.00 |
| Centro | 1 | 0.85 | 164 | 6.10 |
| Rio Comprido | 1 | 0.80 | 162 | 6.17 |
| Botafogo | 0 | 0.99 | 269 | 0.00 |
| Copacabana | 0 | 1.00 | 395 | 0.00 |
| Lagoa | 0 | 1.00 | 284 | 0.00 |
| Sao Cristova | 0 | 0.76 | 244 | 0.00 |
| Tijuca | 1 | 0.95 | 228 | 4.39 |
| Vila Isabel | 0 | 0.93 | 205 | 0.00 |
| Ramos | 0 | 0.78 | 265 | 0.00 |
| Penha & Vigario Geral | 1 | 0.76 | 244 | 4.10 |
| Inhauma | 1 | 0.77 | 216 | 4.63 |
| Meier | 3 | 0.85 | 189 | 15.87 |
| Iraja | 4 | 0.79 | 215 | 18.60 |
| Madureira | 2 | 0.77 | 233 | 8.58 |
| Jacarepagua | 3 | 0.82 | 201 | 14.93 |
| Bangu | 7 | 0.72 | 1097 | 6.38 |
| Campo grande | 5 | 0.73 | 1733 | 2.89 |
| Santa Cruz | 1 | 0.67 | 570 | 1.75 |
| Ilha do Governador | 2 | 0.85 | 368 | 5.43 |
| Anchieta | 4 | 0.72 | 418 | 9.57 |
| Santa Teresa | 0 | 0.83 | 306 | 0.00 |
| Barra de Tijuca | 1 | 1.00 | 518 | 1.93 |
| Pavuna | 4 | 0.69 | 303 | 13.20 |
| Guaritiba | 1 | 0.68 | 259 | 3.86 |
| Rocinha | 0 | 0.67 | 390 | 0.00 |
| Jacarezinho | 0 | 0.64 | 278 | 0.00 |
| Complexo de Alemao | 1 | 0.64 | 293 | 3.41 |
| Mare | 1 | 0.65 | 652 | 1.53 |
| Realengo | 3 | 0.75 | 641 | 4.68 |
| Cidade de Deus | 1 | 0.66 | 328 | 3.05 |

**Appendix 3.**
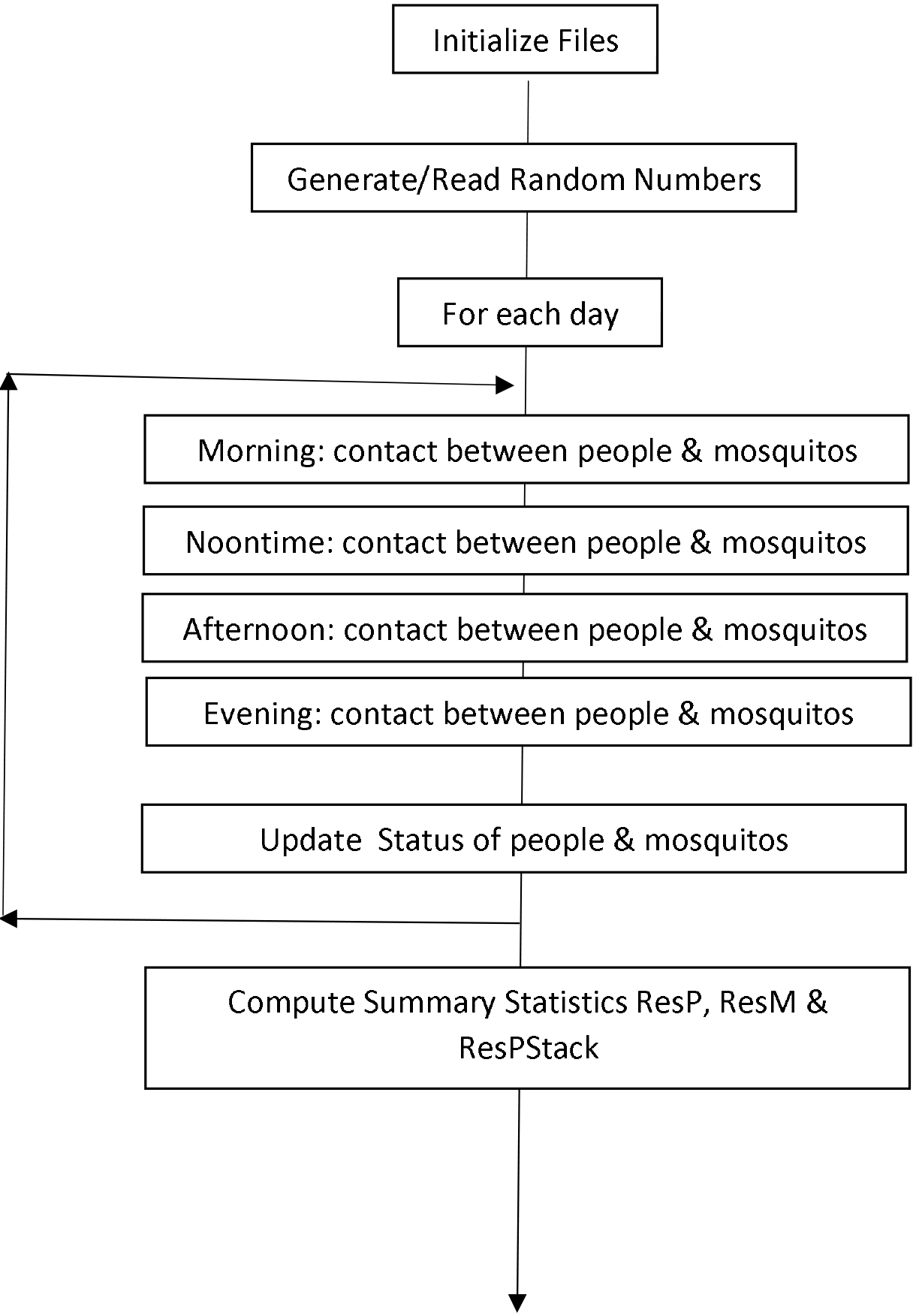
Flow Chart for Agent-based Simulations.

## Acknowledgements

- Health Secretariat of the City of Rio de Janeiro for providing the data;
- Former civil engineer, C Leal, who had worked as an intern in designing the metro system in the Zona Sul, for his insights into the transport system;
- Participants of the Seminar on InfoDengue held at FGV on 19 October 2016, where a preliminary version of the work was presented, for the constructive comments.

